# Norwegian Kveik brewing yeasts are adapted to higher temperatures and produce fewer off-flavours under heat stress than commercial *Saccharomyces cerevisiae* American Ale yeast

**DOI:** 10.1101/2021.06.15.448505

**Authors:** Dimitri Kits, Lars Marius Garshol

**Affiliations:** Independent researcher

**Keywords:** Kveik, brewer’s yeast, *Saccharomyces cerevisiae*, American Ale, temperature, off-flavors

## Abstract

Norwegian kveik are a recently described family of domesticated *Saccharomyces cerevisiae* brewing yeasts used by farmhouse brewers in western Norway for generations to produce traditional Norwegian farmhouse ale. Kveik ale yeasts have been domesticated by farmhouse brewers through serial repitching of the yeast in warm wort (>30°C) punctuated by long periods of dry storage. Kveik yeasts are alcohol tolerant, flocculant, capable of utilizing maltose/maltotriose, phenolic off flavour negative, and exhibit elevated thermotolerance when compared to other modern brewer’s yeasts belonging to the ‘Beer 1’ clade. However, the optimal fermentation and growth temperatures (T_opt_) for kveik ale yeasts and the influence of fermentation temperature of the production of flavour-active metabolites like fusel alcohols and sulfur compounds (H_2_S, SO_2_) are not known. Here we show that kveik ale yeasts have an elevated optimal fermentation temperature (T_opt_) when compared to commercial American Ale yeast (SafAle™ US-05) and that they produce fewer off-flavours at high temperatures (>30°C) when compared to commercial American Ale yeasts. The tested kveik yeasts show significantly higher maximum fermentation rates than American Ale yeast not only at elevated temperatures (>30°C), but also at ‘typical’ ale fermentation temperatures (20°C-25°C). Finally, we demonstrate that kveik ale yeasts are heterogeneous in their T_opt_ and that they attenuate standard wort robustly above their T_opt_ unlike our control American Ale yeast which showed very poor apparent attenuation in our standard wort at temperatures ≫ T_opt_. Our results provide further support that kveik yeasts may possess favourable fermentation kinetics and sensory properties compared to American Ale yeasts. The observations here provide a roadmap for brewers to fine tune their commercial fermentations using kveik ale yeasts for optimal performance and/or flavour impact.

## Introduction

Norwegian kveik ale yeasts are traditional yeast cultures that have been used by farmhouse brewers for the production of beer in western Norway for generations. Kveik are hybrid *Saccharomyces cerevisiae* yeasts that are genetically separate from other ale yeasts, with one haplotype forming a distinct group within the ‘Beer 1’ clade (American, British, German Ale yeasts) and the other forming a distinct group between the ‘Asia’ and ‘Mixed’ groups ^1^. Due to continuous reuse over generations by farmhouse brewers, kveik yeasts show typical phenotypic traits associated with highly domesticated yeasts: high flocculation, ability to utilize maltose/maltotriose, absence of phenolic off flavour (POF-), high ethanol tolerance, and high attenuation ^1^.

Traditional brewing practices in western Norwegian farmhouse brewing, particularly in respect to yeast propagation and management, may have been one factor in the long-term adaptation and evolution in kveik yeast for several distinct phenotypic qualities that have immense relevance for commercial brewers. Farmhouse brewers brewed much less frequently than commercial brewers (typically 1-4 times per year) and kveik ale yeasts were routinely stored dry between use, often for prolonged periods up to 1 year, and not propagated in liquid medium serially prior to pitching ^2^. Kveik yeast are also routinely pitched into wort with very high original extract (>17°P) at high temperatures (25°C-40°C). The most common pitch temperature for Norwegian farmhouse brewers that used kveik was 35-40°C, well above the temperatures typically used for commercial American, British, and German Ales (18-28°C)^3^. The high pitch temperatures and rapid fermentation also resulted in very short fermentation times, with the majority (>65%) of brewers using farmhouse yeast in Norway reporting fermentation times shorter than 50 hours ^2^.

Recent studies characterizing the genetics and brewing characteristics of these traditional kveik ale yeasts clearly demonstrated that these yeasts have high thermotolerance compared to control ale strains ^1^. For example, 19/25 kveik strains were able to grow at 40°C while the control American Ale strain (WLP001) was not. Growth was still evident at 43°C for 19/25 kveik ale strains while only the Belgian Ale strain (WLP570, a member of the ‘Beer 2’ clade) showed growth at this temperature ^1^. All 25 tested kveik yeasts also outperformed American Ale yeast (WLP001) at a fermentation temperature of 30°C when CO_2_ production was measured 24 hours post pitch ^1^.

Though the thermotolerance in kveik ale yeasts is now well described, several phenotypic characteristics have not been investigated. The optimum temperature for growth and fermentation (T_opt_) have not been described for kveik ale yeasts. It is, for example, not understood whether all *S. cerevisiae* yeast that belong to the kveik group have elevated thermal tolerance or whether there is heterogeneity in fermentation performance at different temperatures. Heterogeneity in T_max_ and T_opt_ for kveik ale yeast may be expected, because there is significant heterogeneity known among brewing practices for farmhouse brewers in Norway; pitch temperatures and fermentation times varied and may have resulted in variable adaptations of kveik yeasts to elevated temperatures^3^. It is also not known whether kveik yeasts have a significantly different T_opt_ when compared to commercial American Ale yeasts. Prolonged and distinct propagation of kveik yeasts and commercial brewer’s ale yeast at different temperatures may have significant consequences on the ability of the commercial ale yeasts to tolerate high temperatures. Finally, the influence of elevated fermentation temperatures (> T_opt_, >30°C) on the production of flavour active metabolites for kveik ale yeasts has not been previously investigated.

To address these questions, we performed fermentation tests using a standard wort with four different *Saccharomyces cerevisiae* yeasts – one American Ale yeast (SafAle™ US-05), and three kveik yeasts (Escarpment Labs Laerdal kveik, Lalbrew Voss™ kveik, and Omega Yeast Labs Lutra™ kveik) – at eight different temperatures spanning 20°C-42°C. We analyze the fermentation kinetics of the four ale yeasts as a function of fermentation temperature, determine the T_max_ and T_opt_ of the four strains, and determine the influence of fermentation temperature on the apparent attenuation and sensory characteristics of the finished beer.

## Materials and Methods

### Yeast strains

Four distinct brewer’s yeasts were cultivated and analyzed in this study: a modern commercial ale brewer’s yeast (SafAle™ US-05, Lesaffre) and three distinct Norwegian kveik yeasts – Voss™ (Lesaffre), Lutra™ (Omega Yeast), and Laerdal (Escarpment Labs). SafAle™ US-05 and Lalbrew Voss™ were purchased in dehydrated form, while Lutra™ and Laerdal were purchased in liquid pouch format. The dried yeasts were rehydrated in sterile wort (see Wort preparation) for 30 minutes at room temperature without shaking and immediately streaked onto solid MYGP media (0.3% malt extract, 0.5% peptone, 0.3% yeast extract, 1% dextrose, prepared in tap drinking water, solidified with 1.5% agar) to obtain single isolated colonies. To obtain isolated colonies from Lutra™ and Laerdal, we streaked the liquid yeast (prepared by the manufacturer) directly onto solid MYGP medium.

### Wort preparation

Wort for continuous propagation and fermentation for all yeasts was prepared by extracting and converting the sugars from a 60-minute mash consisting of 60% Pilsner malt, 38% Vienna malt, and 2% Munich malt (Plohberger Malz, Grieskirchen). The 60-minute mash was performed at 65°C with continuous agitation and heating to maintain temperature ±0.5°C. Mash pH was maintained at 5.45-5.5 using a pH meter (LIUMY). Complete conversion after 60 minutes was verified by a refractomer (0-20% Brix ATC), with final densities ranging from 12.1 to 12.6°Plato. A 1-minute incubation at 78°C followed the standard mash to decrease the viscosity of the wort prior to wort sterilization by boiling. Wort was boiled at 100°C for 60 minutes, with a Hallertau Magnum (Mashcamp, Austria) addition at 60 minutes (23 IBU) and a Halltertau Mittelfrueh (Mashcamp, Austria) addition at flameout (0.7 IBU). 0.14 g/L yeast nutrient (Wyeast) was added after 50 minutes of boiling. The sweet wort was then chilled using a stainless-steel immersion coil chiller (Speidel, Ofterdingen, Germany) to yeast pitching temperature. The cooled, sweet wort was aerated for 4 minutes using an aquarium air pump, heat sterilized silicone tubing, and a 5 mm stainless steel airstone. For routine propagation of single isolated colonies in small volumes of wort (<2 L), prior to experiments, we sterilized the bitter wort after mashing using an Instant pot (Instant Pot Duo Nova) using the ‘pressure cook function’. Sterilized wort was transferred under a flame to 50 mL sterile glass bottles, 500 mL sterile borosilicate glass bottles (neoLab, Amazon) or 2 L sterile borosilicate glass bottles (neoLab, Amazon). For this routine propagation, wort was aerated after sterile transfer by shaking the glass vessel.

### Propagation and fermentation

Single isolated colonies of the 4 yeast strains were individually picked from fresh MYGP agar and transferred to 20 mL of sterile wort (see wort preparation) and incubated at 25°C for 48 hours. These 20 mL starters were then each transferred to 200 mL of sterile wort and incubated at 25°C for 48 hours. For each yeast, the 200 mL secondary starter was then used as inoculum for 1 L of sterile wort, which was then incubated at 25°C for 48 hours. The 1 L starters were stored at 4°C for no longer than 2 days prior to pitching for a full scale experiment. HDPE fermenters (15L, Speidel, Ofterdingen, Germany) containing 10 L of wort (see *Wort preparation*) that was prepared by conventional boiling, hopping, cooling, and aeration rather than Instant Pot sterilization (due to volume limitation of the Instant Pot) were inoculated with 1 L of a starter, in triplicate. Triplicate HDPE fermenters were inoculated with each yeast and incubated at 20°C, 25°C, 28°C, 30°C, 33.5°C, 37°C, 40°C, and 42°C in thermally insulated incubators. Temperature in the incubators was controlled by a Temperature controller (Inkbird ITC-308) coupled to a heating mat (21W, Lerway) and refrigerator socket. To measure the temperature, we inserted a temperature probe (Inkbird ITC-308) into a thermowell in the HDPE fermenters. Temperatures were set as mentioned above and controlled to ±0.3°C throughout the fermentation. Fermentation performance was tracked over 144 hours for all strains at all temperatures using a hydrometer (0-20°Plato, Mashcamp, Austria); samples were cooled, and density was already measured at 20°C. 24 to 48 hours after stable final density, each triplicate (per temperature, per yeast strain) was pooled and transferred a Cornelius stainless steel keg. The headspace of each keg was purged 4 times by pressurizing the vessel to 2 bar with 99.9% CO_2_ and releasing the pressure release valve to 1 bar. Each vessel was then carbonated to 2.5 volumes CO_2_ at 4°C for 7 days prior to sensory analysis.

### Sensory analysis

Due to COVID-19 restrictions, a tasting panel consisting of 2 members sampled the finished fermentations. The panel was blind to the identity of the yeast and temperature used for each sample and rated the perceived intensity of 11 different sensory criteria (solvent/fusel, astringent, bitterness, body, floral, fruity, linger, malty, acid, spicy, sulfur) on a scale of 0 (absent) to 5 (severe).

## Results and Discussion

### Kveik yeasts exhibit fast fermentation kinetics within a broad temperature range

SafAle™ US-05 (American Ale) is a commercial brewer’s ale yeast (*Saccharomyces cerevisiae*) used frequently in the production of clean and well-balanced ales. US-05 is closely related to many other dry yeast strains used to produce Pale American ale, IPA, Amber and Brown ales that likely originate from a popular ‘Chico’ strain, which arose in the UK ^4^. These American Ale strains are highly domesticated and possess a large number of desirable qualities for the commercial production of fermented cereal beverages. US-05 shows moderately high attenuation (78-82%), has a high alcohol tolerance (9-11%), tends to stay in suspension well during active fermentation (moderate flocculation), has a wide fermentation temperature range (18-28°C), produces low amounts of esters, and is altogether a robust modern commercial brewer’s yeast.

As a benchmark for performance and reliability, we measured the fermentation kinetics of this commercially used US-05 yeast in a standard wort (12.1-12.5°P) at 8 different temperatures (20°C, 25°C, 28°C, 30°C, 33.5°C, 37°C, 40°C, and 42°C, Figure 1). US-05 exhibited a wide temperature range for fermentation; robust activity was apparent between 20°C and 33.5°C. However, temperatures at or above 37°C completely inhibited fermentation and growth in US-05 (Figure 1). Surprisingly, temperatures within the permissible range for growth (20°C-33.5°C) did not significantly influence the total fermentation time, with temperatures from 20°C to 30°C all reaching final specific gravity (1.010) at 120 hours (5 days) (Figure 1). However, the total fermentation time was extended by 24 hours (144 hours total) at 33.5°C. The maximum temperature for growth (T_max_) of US-05 has not been previously reported to our knowledge. The manufacturer’s recommended temperature range extends to 28°C and previous work has demonstrated that American Ale yeast similar to US-05 (WLP001) grows well at 30°C but not at 40°C ^1^. The T_max_ for other *Saccharomyces cerevisiae* brewing yeasts which have been examined are in the range of 37.5°C −38.5°C ^5^. Our results demonstrate that the T_max_ of US-05 brewers’ dry yeast is 37.0°C and comparable to other top fermenting brewing strains ^5^.

**Figure 1:**
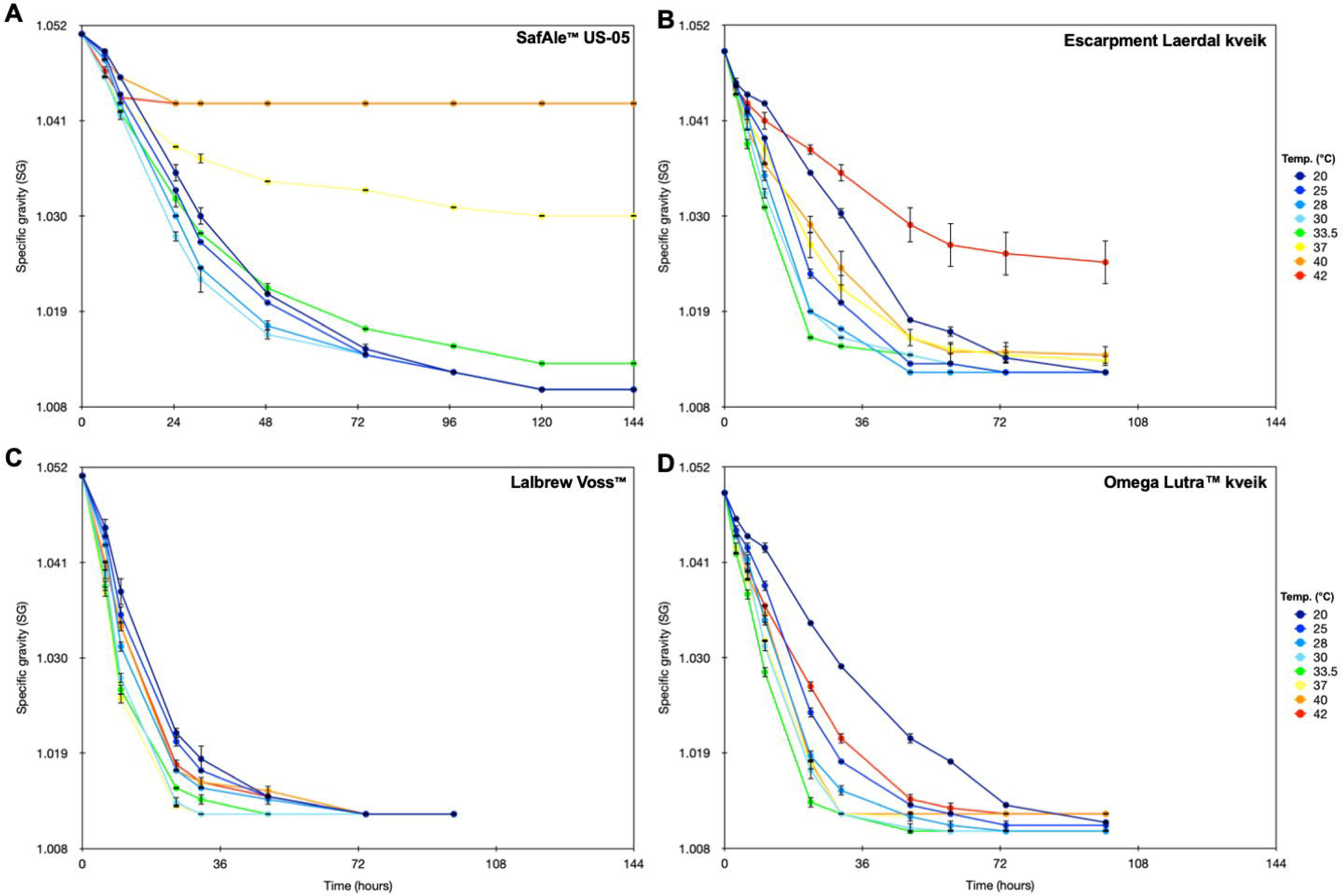
Influence of temperature on the general fermentation performance of four tested *S. cerevisiae* brewer’s yeast strains. Incubations were done in triplicate (n=3) for each temperature for each strain and specific gravity was measured over time as described in the materials and methods. The four panels depict the fermentation kinetics of the four different strains tested: panel A – SafAle™ US-05, panel B – Escarpment labs Laerdal kveik, Panel C – Lalbrew Voss™, Panel D – Omega Yeast labs Lutra™ kveik. Each point represents the mean of three replicates and error bars correspond to the standard deviation of the mean.

Using the same standard wort and temperature range, we next compared the performance and optimal fermentation temperature range of three strains of commercially available, single isolate kveik yeast to that of American Ale yeast. Escarpment Laerdal kveik, Lalbrew Voss™, and Omega Lutra™ were all able to grow at temperatures ranging from 20°C to 42°C and all showed a wider fermentation temperature range than US-05. Consistent with previous reports^1^, the three tested kveik strains showed highly elevated thermal tolerance compared to US-05; all of the three tested kveik yeast were able to grow at 40 and 42°C, while American ale yeast could not grow at temperatures >37°C (Figure 1). In contrast to US-05, fermentation temperature had a significant impact on the total fermentation time for the kveik yeast. Total fermentation time (hours required to reach final specific gravity) ranged from 30 hours (Lalbrew Voss™ at 37°C) to 99.5 hours (Escarpment Laerdal at 20°C and 37°C, Figure 1). Compared to the lowest tested temperature (20°C), all three kveik strains showed markedly decreased fermentation times at elevated temperatures (>20°C); optimal fermentation performance (i.e., short duration of fermentation) was 28°C for Escarpment Laerdal kveik, 37°C for Lalbrew Voss™, and 33.5°C for Omega Lutra™. Very short and rapid (<48 hours) fermentations near the optimum fermentation temperature (T_opt_) are consistent with reported fermentation times for Norwegian farmhouse brewers that use kveik; 75% of farmhouse brewers in Norway reported total fermentation times of <48 hours 2. Generally, warmer temperatures resulted in significantly shorter fermentations up to 40°C, above which all of the tested kveik strains showed extended fermentation times. Despite extended fermentation times for the three kveik strains at temperatures <28°C or > 40°C, the longest total fermentation time for any of the kveik strains (99.5 hours at 20°C for Omega Lutra™ and Escarpment Laerdal) were still significantly shorter than the shortest fermentation time when using US-05 (120 hours, Figure 1). The ability to shorten commercial beer fermentation time has a significant and direct impact on the bottom line of a brewery, since it allows more product to be made per unit of time.

The three tested kveik ale yeasts are single strain isolates that originate from complex mixed cultures, which are usually comprised of several strains of *S. cerevisiae* and sometimes bacteria; Lalbrew Voss™ originates from Sigmund Gjernes Voss kveik, Lutra™ was isolated from Hornindal kveik, and Escarpment Laerdal was isolated from Lærdal kveik. Thus, the fermentation kinetics observed in this study may not be reflective of the complex farmhouse cultures. The individual farmhouse cultures are also distinct from one another, genetically and phenotypically; the recommended pitch temperatures, attenuation, time to harvest, cropping, number of strains present, and other characteristics are distinct and may be a result of the adaptation of each culture to specific brewing practices of the ‘owner’^6^. The most notable difference in the three original farmhouse kveik from which the three isolates were purified is the recommended pitch temperature. The recommended pitch temperature is 30°C for the Hornindal kveik and Laerdal kveik, and 39°C for Sigmund Gjernes Voss kveik ^6^.

### Kveik yeasts demonstrate an elevated temperature optimum compared to modern commercial brewer’s yeast

We define the optimum temperature (T_opt_) as the temperature where the fermentation rate is maximized. Fermentation rate was chosen as the measure, rather than growth rate, because determination of T_opt_ by measuring growth rates is very time consuming and laborious. To determine the T_opt_ for kveik and one American Ale yeast (US-05), we determined the maximum fermentation rates at each temperature for each yeast by calculating the rate of specific gravity decline throughout fermentation (i.e., rate of sugar consumption, Figure 2) and selected the maximum rate as the T_opt_. The T_opt_ of US-05 was 28°C (2.08 specific gravity points consumed per minute ×10^^−5^). This optimum fermentation temperature was slightly lower than previously reported values for non-kveik *S. cerevisiae* brewing yeasts (31.0°C) ^5^. All three kveik strains demonstrated significantly higher thermal fermentation optima when compared to the American Ale yeast; the T_opt_ was 33.5°C, 37°C, and 33.5°C for Escarpment Laerdal, Lalbrew Voss™, and Omega Lutra™, respectively (Figure 2). The higher T_opt_ of Lalbrew Voss™ is also reflected in the higher recommended pitch temperature for the farmhouse culture ^6^.

**Figure 2:**
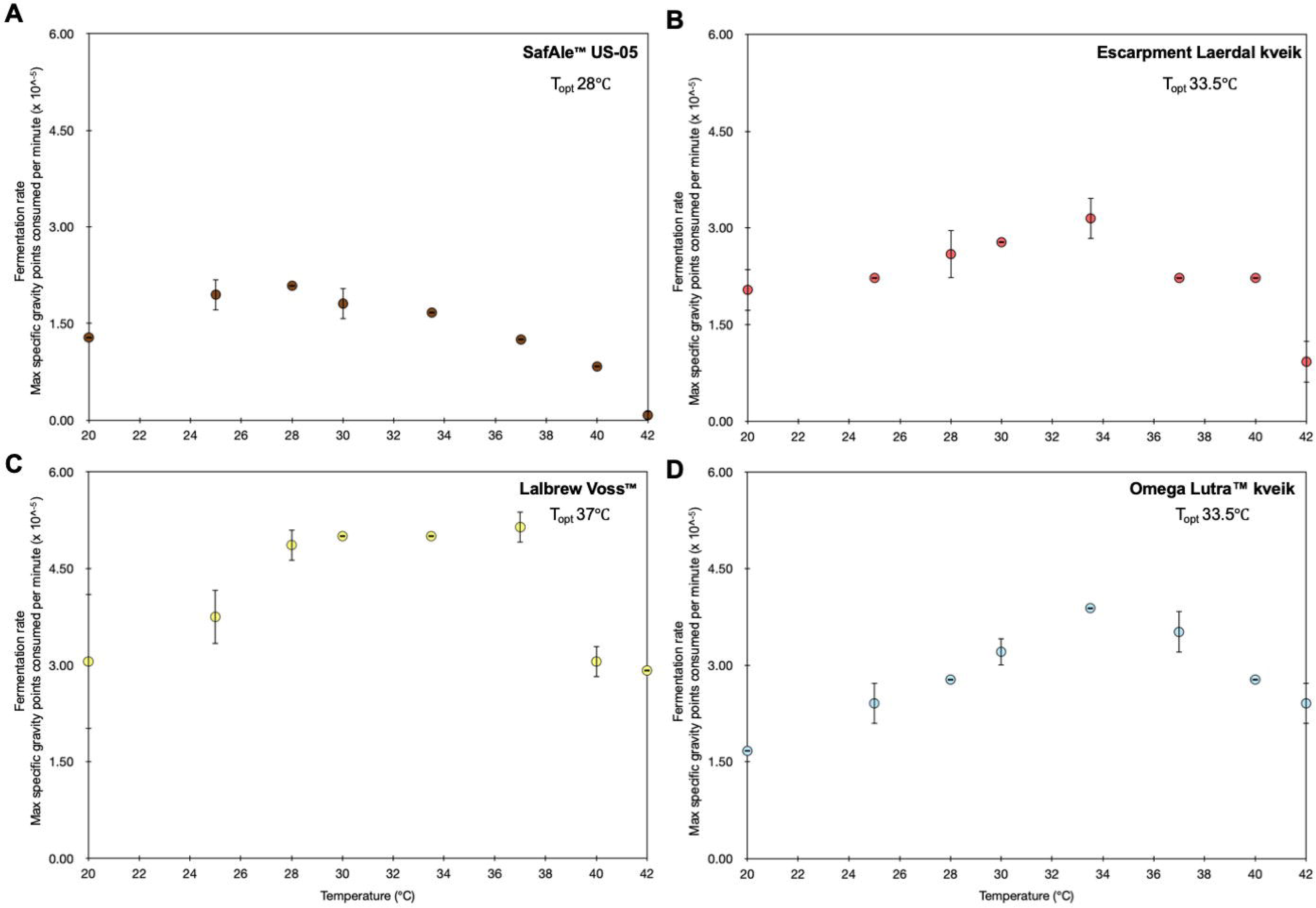
Maximum rates of fermentation based on rates of specific gravity decline per minute for four brewers *S. cerevisiae* strains at eight temperatures ranging from 20°C to 42°C. The four panels depict the fermentation kinetics of the four different strains tested: panel A – SafAle™ US-05, panel B – Escarpment labs Laerdal kveik, Panel C – Lalbrew Voss™, Panel D – Omega Yeast labs Lutra™ kveik. Each point represents the mean of three replicates and error bars correspond to the standard deviation of the mean. The maximum specific gravity points consumption per minute (×10^^−5^) were calculated at each time interval by subtracting the specific gravity at that interval from the previous interval, dividing by the number of minutes elapsed, and then multiplying by 10000; the reported values are absolute values of the actual rates of fermentation, which are negative values (i.e., higher numbers correspond to faster fermentation rates).

There is a strong correlation between wild yeast taxa and their temperature preference^7^. Isolated wild *S. cerevisiae* yeasts have a T_opt_ around ~31.5-35°C and are much more frequently isolated at high temperatures (>20°C) than *S. paradoxus* or *S. eubayanus*, which generally have a T_opt_ significantly lower than *S. cerevisiae* ^8,9^. The measured T_opt_ of the three kveik strains we tested (33.5-37.0°C) was similar to the range reported for wild *S. cerevisiae* yeasts but significantly higher than the T_opt_ of the commercial American Ale yeast (28°C). The difference in T_opt_ between kveik ale yeasts and commercial brewer’s yeasts may be due to differences in yeast handling over time between traditional farmhouse brewing and modern commercial beer production. Norwegian farmhouse brewers pitched yeast serially into warm wort (>28°C) while modern American Ale yeast is serially repitched for an extended number of brewing cycles in cool wort (<20°C). It is possible that commercial ale yeast lost the ability to tolerate high temperatures as a consequence of long-term adaptation to commercial brewing processes, while the traditional farmhouse brewing practice of pitching dry yeast into warm wort may have maintained high thermal tolerance and high thermal optima in kveik ale yeasts^10^.

The maximum rate of fermentation per unit time, as measured by the maximum rate of wort specific gravity decline per minute, is also considered to be a measure of the productivity of the yeast at a particular temperature. Comparing fermentation rates between the tested yeast strains at various temperatures not only describes the differences in T_opt_ between the different strains, it also allows a direct comparison of fermentation speed at a particular temperature (Figure 5). At every temperature we tested, all three tested kveik yeasts exhibited a maximum fermentation rate that was faster than that of US-05. For example, the maximum fermentation rate at 20°C was 2.38-fold higher (3.06 vs 12.8) for Voss kveik than that observed for US-05 (Figure 5). This trend was also apparent at the respective optimum fermentation temperatures for US-05 and Voss – the maximum fermentation rate at 37°C for Voss kveik was 2.47-fold higher (5.14) than the rate observed for US-05 at 28°C (2.08, Figure 5). This demonstrates that kveik yeasts not only have higher fermentation T_opt_ but that they also have a significantly higher substrate turnover when compared to commercial brewer’s yeast at comparable temperatures. The broader fermentation temperature range and higher substrate turnover rates are tremendously important in the commercial brewing sector, where these unique properties enable energy savings in wort cooling and lead to increase in brewery product output due to shorter fermentation times.

An alternative way to assess activity of the tested yeast strains is the total fermentation time rather than the maximum rate of fermentation per unit time. The maximum rate is reflective of the peak in metabolic activity (i.e., maximum velocity), while the total fermentation time is also influenced by how long a particular level of fermentation is sustained and also the ability of each strain to consume maltose/maltotriose (i.e., enzyme affinity). For the kveik strains, the shortest total fermentation time (hours required to reach final specific gravity) for each strain was generally observed at or near the T_opt_. The shortest total fermentation time was at an incubation temperature of 37.0°C for Voss kveik (T_opt_ 37.0°C), 30-33.5°C for Lutra kveik (T_opt_ of 33.5°C), and deviated slightly only for Laerdal kveik, for which the shortest fermentation was at 28°C but the T_opt_ was 33.5°C. The total fermentation time for US-05 was not strongly influenced by temperature (120 hours at 20-30°C and 144 hours at 33.5°C). The weak influence of fermentation temperature on total fermentation time in US-05 seems due to the very weak stimulation of the maximum rate of fermentation by temperature; the maximum rate at T_opt_ was only ~1.6 fold higher than at the lowest tested temperature (20°C). So though the total fermentation time and the maximum rate of fermentation are different measures, the two largely agreed across our data set.

### Kveik yeasts show robust attenuation in wort at elevated temperatures

The apparent attenuation describes the extent to which a wort is fermented by yeast and can be calculated by comparing the terminal specific gravity of the green beer to the original specific gravity of the bitter wort. The degree to which a particular yeast strain attenuates a wort is dependent on fermentation conditions such as temperature, pitch rate, wort osmolarity, yeast health, and mashing regime, but is also highly strain dependent ^11^. We compared the apparent attenuation of the three kveik strains to the commercial ale yeast US-05 as a product of various fermentation temperatures. Higher temperature fermentations are known to generally decrease the apparent attenuation of yeast in wort due to temperature stress ^12^. US-05 showed an apparent attenuation of 80% when fermented within or near the manufacturer’s recommended temperature range (20°C-28°C, Figure 3). A significant decrease in the apparent attenuation was evident at fermentation temperatures >30°C, with apparent attenuation levels of 77.4% at 33.5°C, and only 40% at T_max_ of 37°C (Figure 3). For US-05, the apparent attenuation matched the temperature performance curve (Figure 2, Panel A) where full expected apparent attenuation was reached only up to 2°C above T_opt_ (28°C for US-05).

**Figure 3:**
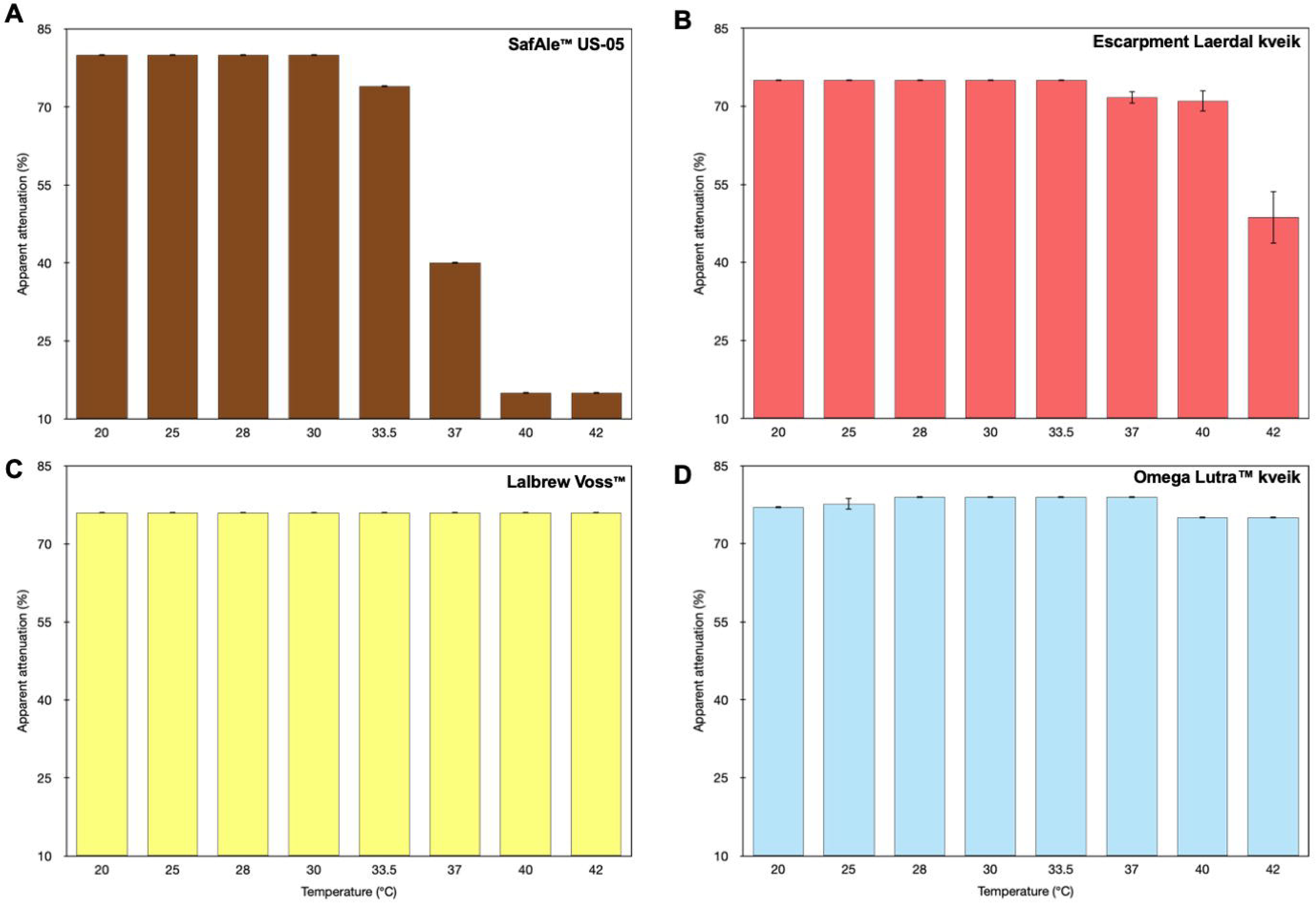
Influence of temperature on the apparent attenuation (%) of fermentations in standard wort for four brewer’s *S. cerevisiae* strains. The four panels depict the fermentation kinetics of the four different strains tested: panel A – SafAle™ US-05, panel B – Escarpment labs Laerdal kveik, Panel C – Lalbrew Voss™, Panel D – Omega Yeast labs Lutra™ kveik. Each point represents the mean of three replicates and error bars correspond to the standard deviation of the mean. Apparent attenuation was calculated by subtracting the residual extract (final gravity) from the initial extract (original gravity) and dividing by 1 subtracted from the initial extract and reported as a percentage.

Kveik ale yeasts have previously been demonstrated to attenuate wort in a broad range (60-90%), allowing commercial brewers to select strains suitable for a preferred attenuation target ^1^. However, how temperature influences apparent attenuation by kveik yeasts is unknown. All three kveik strains we tested showed more consistent apparent attenuation at a broader temperature range than US-05 (Figure 3). Escarpment Laerdal kveik is mildly attenuative (75% apparent attenuation) at temperatures at or below its T_opt_ (33.5°C or lower) and undergoes a small decrease in the apparent attenuation (71% apparent attenuation) at temperatures up to 40°C (Figure 3). The apparent attenuation of Omega Yeast Labs Lutra™ kveik in standard wort is also robust throughout the tested temperature range; we observed 77-79% apparent attenuation from 20°C to 37°C with a small decline (75% apparent attenuation) at 40-42°C (Figure 3). Lalbrew Voss™ kveik was the most consistent and robust in terms of apparent attenuation; the apparent attenuation in standard wort was 77% within the entire tested temperature range (20-42°C). The range in observed attenuative capabilities of the tested kveik strains also correlates with the recommended pitching temperature for the original farmhouse cultures; Lalbrew Voss™ was isolated from Sigmund Gjernes Voss kveik, which has a significantly higher recommended pitching temperature (39°C) than the farmhouse cultures where Lutra™ and Laerdal were isolated from (Hornindal kveik and Lærdal kveik, recommended pitch temperature of 30°C)^6^. The ability of kveik yeasts to fully attenuate wort at a broader temperature range is another highly desirable and relevant yeast characteristic for commercial brewers as it allows the brewer flexibility in temperature control or flexibility in choosing a particular temperature-driven flavour profile without a decrease in attenuation.

### Kveik yeasts produce less off-flavours at elevated temperatures than modern S. cerevisiae brewer’s yeast

Fermentation of saccharides by yeast in wort results in the production of CO_2_, ethanol, and many other small flavour-active metabolites such as organic acids, aldehydes, esters, sulfur compounds, phenolic compounds, ketones, as well as aromatic higher alcohols known as fusel alcohols ^13^. Flavour contributions from higher alcohols can be positive when they serve as precursors to acetate esters but in high concentrations are considered to impart off-flavours to beer ^14^. Fermentation conditions such as fermentation temperature, wort aeration, wort osmolarity, increased head pressure, micronutrient content, and aging are known to influence higher alcohol production by brewer’s yeast ^15^. In particular, higher temperatures and faster yeast growth rates are known to increase final concentrations of higher alcohols in finished beer ^16^. Temperature stress is also known to increase the concentrations of other known off-flavours in beer, such as volatile sulfur compounds ^17^.

To determine if increased fermentation temperature increases the production of off-flavours such as higher alcohols and sulfur compounds in kveik yeast, we subjected beers fermented with three different kveik ale strains at eight different temperatures to a sensory analysis with a tasting panel that rated the beers quantitatively on 11 different sensory criteria (solvent/fusel, astringent, bitterness, body, floral, fruity, linger, malty, acid, spicy, sulfur)(Figure 4). We then compared these results with that of the US-05 American Ale yeast to test whether the responses to high temperatures different between the commercial ale yeast and domestical farmhouse kveik yeast.

**Figure 4:**
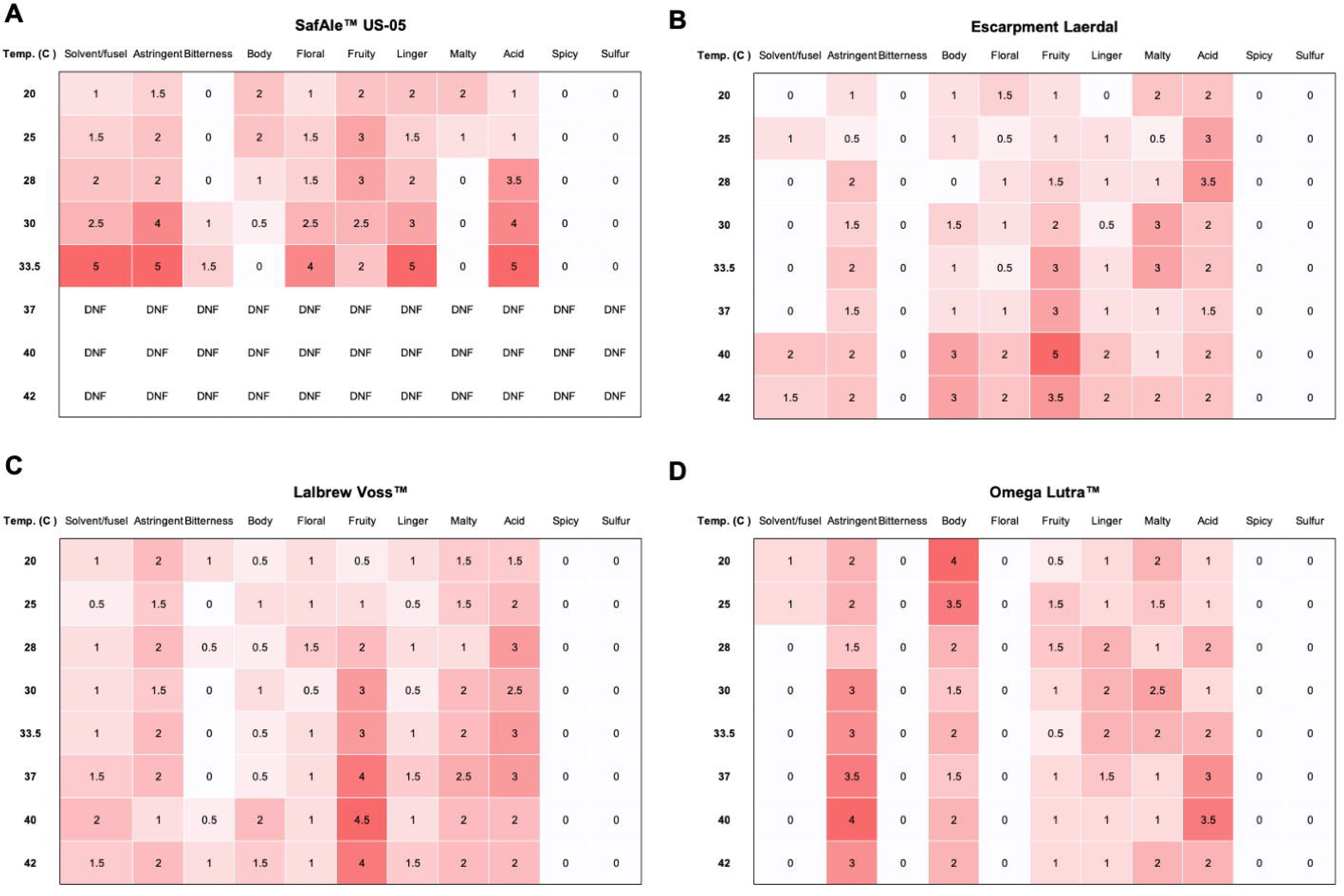
Intensity of 11 quantitative sensory descriptors of bright beer fermented by four brewer’s *S. cerevisiae* strains at eight temperatures ranging from 20°C to 42°C. The four panels depict the fermentation kinetics of the four different strains tested: panel A – SafAle™ US-05, panel B – Escarpment labs Laerdal kveik, Panel C – Lalbrew Voss™, Panel D – Omega Yeast labs Lutra™ kveik. Each tile corresponds to one temperature for one sensory descriptor out of a maximum intensity of 5; tiles are shaded according to the intensity with darker corresponding to a higher intensity. Due to COVID-19 restrictions, a tasting panel consisting of two members conducted the sensory analysis and each tile corresponds to the average of the two values; the tasting panel was blinded to the temperature and strain tested, for further information see the materials and methods.

**Figure 5:**
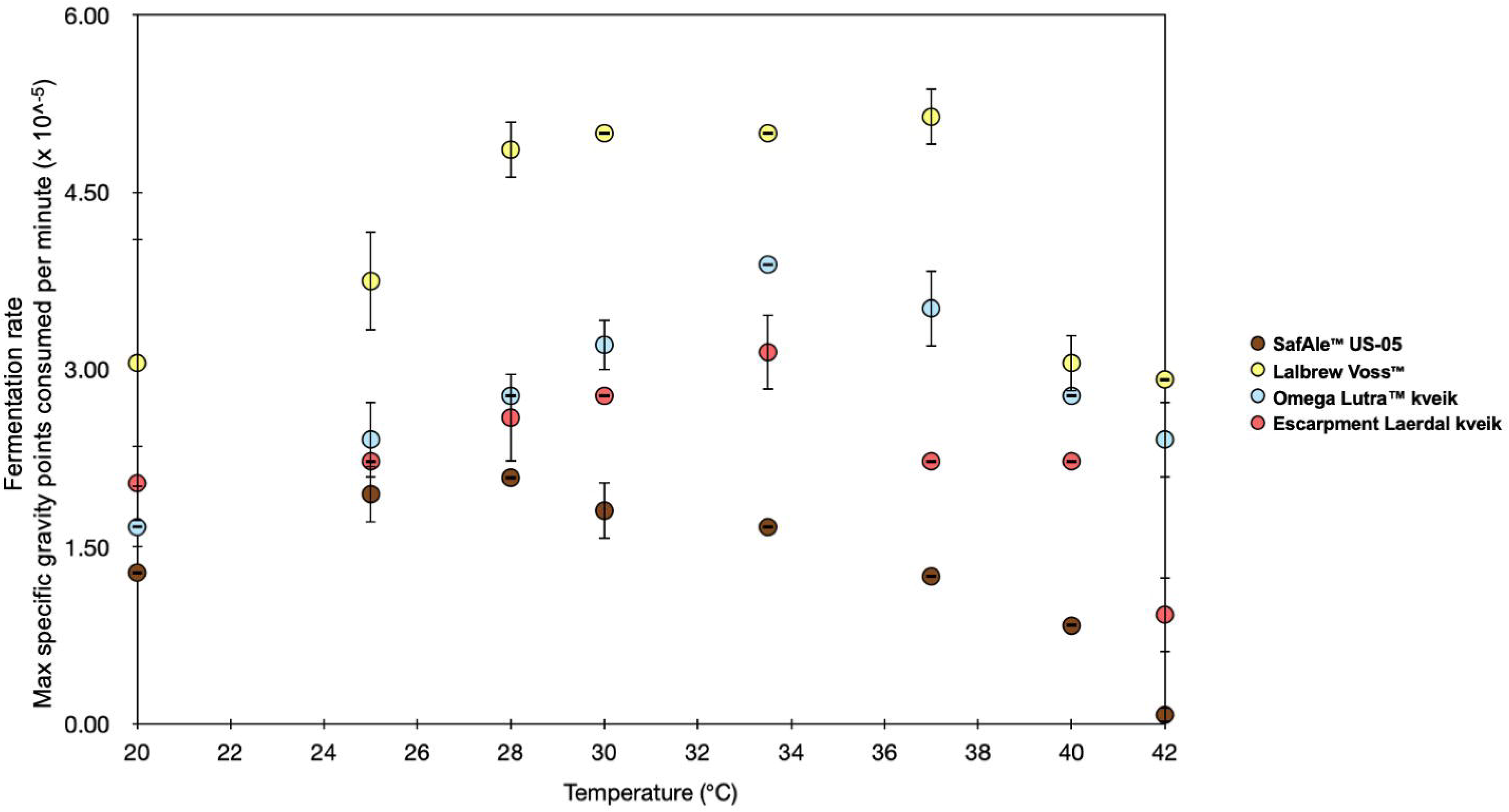
Overlay of maximum rates of fermentation for four brewer’s *S. cerevisiae* strains at eight temperatures ranging from 20°C to 42°C. Each point corresponds to the maximum specific gravity points consumed per minute (×10^^−5^) for four different S. cerevisiae strains; SafAle™ US-05 is depicted in brown, Lalbrew Voss™ kveik is depicted in yellow, Omega Yeast Labs Lutra™ kveik in blue, and Escarpment labs Laerdal kveik in fuchsia. Each point represents the mean of three replicates and error bars correspond to the standard deviation of the mean. The maximum specific gravity points consumption per minute (×10^^−5^) were calculated at each time interval by subtracting the specific gravity at that interval from the previous interval, dividing by the number of minutes elapsed, and then multiplying by 10000; the reported values are absolute values of the actual rates of fermentation, which are negative values (i.e., higher numbers correspond to faster fermentation rates).

At typical ale fermentation temperatures of 20°C to 25°C, standard wort fermented with US-05 exhibited a neutral, clean, and well-balanced flavour profile dominated by light fruit esters, low astringency, low acidity, and light-to-medium body (Figure 4 Panel A). We observed a strong trend of increasing off-flavours – ‘solvent/fusel’, ‘astringent’, ‘linger’ and ‘acid’ – with increasing fermentation temperature. The solvent-like higher alcohols, astringency, and acidity became significantly more pronounced at 30°C, just above the T_opt_ of US-05. At the maximum temperature that allowed a near-normal apparent attenuation (33.5°C, apparent attenuation of 74%), the higher alcohols (‘solvent/fusel’), astringency, and acidity dominated the aroma and flavour of the finished beer (Figure 4 Panel A). Sensory analysis was not possible for beers fermented with US-05 at temperatures >33.5°C due to the extremely high residual extract levels from temperature-dependent stress and growth inhibition.

Sensory analysis of beer fermented with the three tested kveik ale yeasts at 20°C to 25°C demonstrated that each kveik strain has a unique flavour profile and that the three kveik ale yeasts together were distinct from the US-05 American Ale yeast (Figure 4). At these temperatures, the kveik strains were generally perceived as more acidic and more astringent than US-05 but had lower ‘estery’ fruitiness (Figure 4 Panels B-D). At temperatures near the T_opt_ (33.5°C to 37°C), the fruity esters and astringency were mildly elevated for the kveik strains when compared to lower temperatures. These results are congruent with previous work showing that kveik ale yeast produce increased concentrations of fatty acid esters such as ethyl caproate (‘pineapple’), ethyl caprylate (‘tropical fruit’), and ethyl decanoate (‘apple’) at 30°C when compared to American Ale yeast (WLP001) ^1^. The same study also found that some kveik ale yeast also produce lower quantities of fusel alcohols at 30°C. The influence of temperatures above the optimum fermentation temperature on the production of flavour active metabolites produced by kveik ale yeast has not been investigated so far. We demonstrate here that increasing the fermentation temperatures to above T_opt_ (>37°C) for kveik resulted in a further increase in fruit esters but did not increase the perceived astringency or solvent-like higher alcohols (Figure 4 Panels B-D). In some cases (Lalbrew Voss™, Omega Lutra™), the astringency and acidity were perceived to be lower at 40°C-42°C when compared to temperatures near T_opt_ (Figure 4 Panels C-D). It should be noted that due to methodological limitations, this does not mean the metabolites contributing to these flavours are present in lower concentrations; rather, it is possible that the higher abundance of other flavour active compounds (i.e., esters) causes a relative decrease in the taste perception of other compounds without a true decrease in metabolite concentrations. Nevertheless, our results suggest that the excellent thermal tolerance shown by kveik yeasts may allow them to produce lower quantities of off-flavour associated metabolites (especially higher alcohols and sulfur compounds) at very high fermentation temperatures (>37°C). The low quantities of higher alcohols, sulfur compounds, and diacetyl produced by kveik at high temperatures with fast fermentations is another desirable quality for commercial brewing.

### Conclusions

Norwegian kveik ale yeasts are traditional yeasts used and domesticated by farmhouse brewers in western Norway over generations and are genetically and physiologically distinct from modern commercial brewer’s yeast. Kveik ale yeasts have previously shown to be flocculant, thermotolerant, POF-, hybrids that are resistant to high ethanol concentrations and desiccation. We demonstrate here that kveik ale yeasts are heterogeneous in their temperature optimum for fermentation (T_opt_) but also that they have a significantly elevated T_opt_ when compared to commercial American Ale yeast. Further, we show that at least three kveik strains have a very broad fermentation temperature range and show consistent attenuation in wort even at very high temperatures (≫37°C). Finally, using sensory analysis, we provide evidence that kveik yeasts produce fewer off-flavours in beer when fermented at very high temperatures compared to American Ale yeast. In particular, Omega Lutra™ kveik displayed not only exceptional fermentation kinetics and apparent attenuation at a broad temperature but also a restrained ester profile, which makes this yeast particularly suitable for the commercial production of American, British, and German Ales.

## Contributions

D.K. and L.M.G. conceived the study. D.K. performed the experiments. D.K. wrote the study with input from L.M.G.

## References

1. Preiss, R. Tyrawa, C. Krogerus, K. Garshol, L. M. & van der Merwe, G. Traditional Norwegian Kveik Are a Genetically Distinct Group of Domesticated Brewing Yeasts. Front. Microbiol. 9, 2137 (2018).

2. Garshol, L. M. Historical Brewing Techniques: The Lost Art of Farmhouse Brewing. (2020).

3. Garshol, L. M. Pitch Temperatures in Traditional Farmhouse Brewing. Journal of the American Society of Brewing Chemists 79, 181–186 (2020).

4. Large, C. R. L. et al. Genomic stability and adaptation of beer brewing yeasts during serial repitching in the brewery. doi:10.1101/2020.06.26.166157.

5. Walsh, R. M. & Martin, P. A. GROWTH OF SACCHAROMYCES CEREVISIAE AND SACCHAROMYCES UVARUM IN A TEMPERATURE GRADIENT INCUBATOR. Journal of the Institute of Brewing vol. 83 169–172 (1977).

6. Garshol, L. M. The Farmhouse Yeast Registry. MBAA Technical Quarterly vol. 57 123–128 (2020).

7. Sylvester, K. et al. Temperature and host preferences drive the diversification of Saccharomyces and other yeasts: a survey and the discovery of eight new yeast species. FEMS Yeast Res. 15, (2015).

8. Robinson, H. A., Pinharanda, A. & Bensasson, D. Summer temperature can predict the distribution of wild yeast populations. Ecol. Evol. 6, 1236–1250 (2016).

9. Salvadó, Z. et al. Temperature adaptation markedly determines evolution within the genus Saccharomyces. Appl. Environ. Microbiol. 77, 2292–2302 (2011).

10. Garshol, L. M. Pitch temperatures in traditional farmhouse brewing. doi:10.31235/osf.io/wmyfe.

11. Jin, Y.-L. & Alex Speers, R. Flocculation of Saccharomyces cerevisiae. Food Research International vol. 31 421–440 (1998).

12. Gibson, B. R., Lawrence, S. J., Leclaire, J. P. R., Powell, C. D. & Smart, K. A. Yeast responses to stresses associated with industrial brewery handling: Figure 1. FEMS Microbiology Reviews vol. 31 535–569 (2007).

13. Stewart, G. The Production of Secondary Metabolites with Flavour Potential during Brewing and Distilling Wort Fermentations. Fermentation vol. 3 63 (2017).

14. Dufour, J.-P., Malcorps, P. H. & Silcock, P. Control of Ester Synthesis During Brewery Fermentation. Brewing Yeast Fermentation Performance 213–233 (2008) doi:10.1002/9780470696040.ch21.

15. Kunze, W. Technologie Brauer und Mälzer. (1994).

16. Landaud, S. Latrille, E. & Corrieu, G. Top Pressure and Temperature Control the Fusel Alcohol/Ester Ratio through Yeast Growth in Beer Fermentation. Journal of the Institute of Brewing vol. 107 107–117 (2001).

17. Linderholm, A. L., Findleton, C. L., Kumar, G., Hong, Y. & Bisson, L. F. Identification of genes affecting hydrogen sulfide formation in Saccharomyces cerevisiae. Appl. Environ. Microbiol. 74, 1418–1427 (2008).

